# Effect of expression of human glucosylceramidase 2 isoforms on lipid profiles in COS-7 cells

**DOI:** 10.1101/2020.07.06.190314

**Authors:** Peeranat Jatooratthawichot, Chutima Talabnin, Lukana Ngiwsara, Yepy Hardi Rustam, Jisnuson Svasti, Gavin E. Reid, James R. Ketudat Cairns

## Abstract

Glucosylceramide (GlcCer) is a major membrane lipid and the precursor of gangliosides. It is continuously formed and degraded in glycosphingolipid metabolism. GlcCer is mainly degraded by two enzymes, lysosomal acid β-glucosidase (GBA) and nonlysosomal β-glucosidase (GBA2). Deficiencies of GBA and GBA2 affect glycosphingolipid metabolism, resulting in neurological diseases, such as Gaucher Disease and Hereditary Spastic Paraplegia. To understand which GBA2 isoforms are active and how they affect glycosphingolipid levels in cells, we expressed nine human GBA2 isoforms in COS-7 cells, confirmed their expression by qRT-PCR and western blotting, and assayed their activity to hydrolyze 4-methylumbelliferyl-β-D-glucopyranoside (4MUG) in cell extracts. Human GBA2 isoform 1 showed high activity, while the other isoforms had activity similar to the background. Comparison of sphingolipid levels by ultra-high resolution/ accurate mass spectrometry (UHRAMS) analysis showed that isoform 1 overexpression increased ceramide and decreased hexosylceramide levels compared to control and other isoforms. Comparison of ratios of glucosylceramides to the corresponding ceramides in the extracts indicated that GBA2 isoform 1 has broad specificity for the lipid component of glucosylceramide. These studies suggest that only one GBA2 isoform 1 is active and affects sphingolipid levels in the cell, acting on glucosylceramides with a wide range of lipid components. Our study provides new insights into how increased breakdown of GlcCer affects cellular lipid metabolic networks.

## Introduction

Sphingolipid metabolism affects many processes in cellular biology, including apoptosis, cell-cycle arrest, differentiation, migration, proliferation, and senescence (1). In addition, sphingolipids contribute to cell signaling as an important component of cellular membrane, where they help maintain the integrity of membrane structure and organization (2). Sphingolipids have been implicated in metabolism, neurodevelopment, inflammation, cancer, and several other physiological and pathological processes (3–6).

*De novo* synthesis of sphingolipids begins in the endoplasmic reticulum (ER) with condensation of serine and palmitoyl-CoA by serine palmitoyltransferase (SPT), followed by reduction of the 3-ketosphingonine to sphingonine, which is converted to ceramide by N-acylation and oxidation to form a double bond (7). Furthermore, ceramide is a precursor for synthesis of hexosylceramide, sphingomyelin, and sphingosine (1). Hexosylceramides, also known as cerebrosides, include glucosylceramide (GlcCer) and galactosylceramide (GalCer), which serve as precursors for synthesis of more complex glycosphingolipids. A portion of the ceramide may be converted to GalCer by galactosylceramide synthase in the ER. The remaining ceramide is transported to the Golgi complex, where one of two enzymes catalyzes the synthesis of two complex sphingolipids, GlcCer and sphingomyelin. Transfer of phosphorylcholine from phosphatidylcholine (PC) to Cer on the lumen side of the Golgi membrane by sphingomyelin synthase (SMS) produces sphingomyelin, while on the cytosolic side of the Golgi membrane, glucosylceramide synthase (GCS) converts ceramide to glucosylceramide through the addition of a glucosyl group from UDP-glucose. GlcCer is then transported to the ER and flipped in the membrane to put the glucosyl head group in the lumen, then transferred to the plasma membrane via the Golgi and trans-Golgi network (6). Furthermore, in the Golgi GlcCer can act as the building block for lactosylceramide and other complex glycosphingolipids, such as gangliosides (8).

GlcCer and its more complex products are ultimately fated to undergo degradation through hydrolytic pathways, mostly in the lysosome (9, 10). GlcCer is degraded by glucosylceramidase (glucocerebrosidase, acid β-glucosidase, GBA, E.C. 3.2.1.45) and glucosylceramidase 2 (nonlysosomal β-glucosidase, GBA2, EC 3. 2. 1. 45) to release ceramide and glucose (7). GBA is stimulated by glycerophospholipids, such as phosphatidylglycerol (PG), phosphatidic acid (PA), phosphatidylinositol (PI), and phosphatidylserine (PS) (6, 11). GBA dysfunction in hydrolyzing GlcCer results in the accumulation of GlcCer, and this GBA deficiency is sometimes triggered as a secondary defect of the accumulation of other lipids, such as cholesterol, and GM1 and GM2 ganglioside (6, 7, 12). GBA dysfunction causes Gaucher disease, which is a common lysosomal storage disorder, resulting mainly in GlcCer accumulation, particularly in macrophages (13, 14). Deficiency of GBA has been implicated in the etiology of Parkinson’ s disease (15). In contrast, the biological significance of GBA2 is less well understood. GBA2-deficient mice present with male infertility (16), but humans carrying mutations in the GBA2 gene are affected with a cerebellar ataxia with spasticity or spastic paraplegia, often associated with thin corpus callosum and cognitive impairment as the disease progresses (SPastic Gait locus #46, SPG46) (17).

Alternative splicing generates different transcripts from the same gene and affects the expression levels, stability, half-life and localization of the RNA messengers (18). It has the potential to generate several protein isoforms with different biological properties, protein– protein interactions, subcellular localization, signaling pathway, or catalytic ability (19). The relevance of alternative splicing is made clear by certain point mutations. For instance, the g. 12599C > A (c. 999 + 242C > A mutation, found deep in intron 7 of the GBA gene, creates donor splice site 3 nucleotides 5’ of this mutation, leading to aberrant splicing that results in the insertion of the first 239nt of intron 7 (20). In contrast, the biological significance of GBA2 alternative splicing in neurological diseases has yet to be reported.

In order to understand the biological significance of GBA2 isoforms on sphingolipid metabolism, we have studied GBA2 isoforms by transient overexpression of isoforms predicted by RNA sequencing in mammalian cells, followed by activity assay with artificial substrate (4MUG) and assessment of lipid changes in the cells, including that of the natural substrate (GlcCer). We have found that GBA2 isoform 1 is the only active isoform showing high activity to hydrolyze 4MUG and its overexpression strongly affected the relative levels of ceramide, hexosylceramide and sphingomyelin.

## MATERIALS AND METHODS

### Reagents

Dulbecco’s Modified Eagle Medium, Penicillin/streptomycin (Pen Strep), trypsin/EDTA, and fetal bovine serum were obtained from Thermo Fisher Scientific (Waltham, MA, USA). The coding sequences for 9 isoforms of human GBA2: major transcript isoform 1 (NM_020944. 3), isoform 2 (NM_001330660. 1), isoform X1 (XM_006716809. 3), isoform X2 (XM_005251526. 4), isoform X3 (XM_017014937. 1), isoform X4 (XM_017014938. 1), isoform X6 (XM_017014940. 1), isoform, X7 (XM_017014941. 1), and isoformX8 (XM_017014942. 2) were synthesized and inserted into the pcDNA3. 1+/c-(k) - dyk expression vector for mammalian cells by GenScript Corporation (Piscataway, NJ USA). The human GBA2 peptide CRRNVIPHDIGDPDD was synthesized and an anti-peptide antibody to it was also generated at Genscript Corp. Anti-β-actin and anti-Flag-tag antibodies were from Cell Signaling Technology (Danvers, MA, USA). Conduritol-β-epoxide and 4-methylumbelliferyl-β-D-glucuronide were from Sigma-Aldrich (St. Louis, MO, USA). Deuterated internal standard lipids phosphatidylcholine (PC 15:0/18:1-d7), phosphatidylethanolamine (PE 15: 0/18: 1-d7), phosphatidylserine (PS 15: 0/18: 1-d7), phosphatidylglycerol (PG 15: 0/18: 1-d7), phosphatidylinositol (PI 15:0/18:1-d7), phosphatidic acid (PA 15:0/18:1-d7), lysophosphatidylcholine (LPC 18:1-d7) lysophosphatidylethanolamine (LPE 18:1-d7), cholesterol ester (18:1-d7), monoacylglycerol (MG 18:1-d7), diacylglycerol (DG 15:0/18:1-d7), triacylglycerol (TG 15:0/18:1-d7/15:0), SM (d18:1/18:1-d9), and Cer (d18:1-d7/15:0) were from Avanti Polar Lipids (Alabaster, AL, USA). Ammonium formate was from Alfa Aesar (Ward Hill, MA, USA). Propanol, methanol, and water were from J.T. Baker (Phillipsburg, NJ, USA) and chloroform was from EMD chemicals (Billerica, MA, USA). All solvents used were high performance liquid chromatography grade, and all lipid extraction and storage solvents contained 0. 01% butylated hydroxytoluene (BHT) from Sigma Aldrich.

### Cell culture and transfection

COS-7 cells (African green monkey kidney) were seeded into 75 cm^2^ cell culture flasks with DMEM containing 10% Pen-Strep and 10% fetal bovine serum. Cell lines were grown in a 5% CO_2_ incubator at 37 °C. One million COS-7 cells were seeded into a 6 cm^2^ plate with DMEM medium, and then the COS-7 cells were cultured to 80-90% confluency overnight. The medium was removed and the plate washed once with sterile 1X PBS, then 2 mL of Opti-MEM (reduced serum medium, Thermo Fischer Scientific) was added into the plate. COS-7 cells were transfected with the cDNA-encoding human GBA2 isoforms in the pcDNA3. 1+/c-(k) - dyk mammalian expression vector with lipofectamine 2000 reagent (Thermo Fischer Scientific), according to manufacturer’ s instructions. After 6 hours, the Opti-MEM was removed, and replaced by DMEM complete medium with 1% Pen Strep. COS-7 cells were incubated for 48 hours or 72 hours, then the cells were washed with PBS, scraped in PBS, and stored at −80 °C until use.

### RNA extraction and Quantitative RT-PCR

COS-7 cells were collected at 48 h and 72 h after transfection. RNA was extracted in Trizol reagent (Thermo Fischer Scientific), according to manufacturer’s instructions. The first stand cDNA was generated by SuperScript™ III Reverse Transcriptase cDNA synthesis Kit (Thermo Fischer Scientific), and the synthesized cDNA synthesis reaction was stored at −20 °C until use. RT-PRC was performed using SYBR green/Rox qPCR master mix (Thermo Fischer Scientific) on the LightCycler® 480 II Instrument (Rosche Molecular Systems, Inc., Pleasanton, CA, USA). The primers for qPCR were GBA2-Forward: CCACTACAGGCGGTATACAA and GBA2-reverse: GATCTGTCATCCAATACCGG, and β-actin-Forward: 5’-GATCAGCAAGCAGGAGTATGACG-3’ and β-actin-reverse: 5’-AAGGGTGTAACGCAACTAAGTCATAG-3’

### Protein collection and Western blotting analysis

The medium was removed and cells washed 2 times with ice-cold 1X PBS, then the COS-7 cells were collected in 1 mL of ice-cold 1X PBS by scraping, and the cell suspension was transferred to a 1. 5 mL tube on ice. Then, the cell suspension in 1X PBS was sonicated on ice. The extracted cells were diluted 1: 20 in 1X PBS and the protein concentration was measured with a Pierce™ BCA Protein Assay kit from Thermo Fischer Scientific (#23225). Proteins were separated by SDS-PAGE using the Criterion system (BioRad) and transferred to nitrocellulose membrane by wet western blotting transfer in 50 mM Tris-base, 40 mM glycine, and 20% methanol. Blots were blocked by 5% skimmed milk in 0.05% PBST for 1 hour, and washed with 0. 05% PBST, then incubated with anti-GBA2 antibody (1: 100), anti-FLAG antibody (1: 1000), or anti-β-actin antibody (1: 2000) as primary antibody overnight. After washing 3 times with PBST, goat anti-rabbit/HRP (Genscript) and rabbit anti-mouse/HRP (DAKO)-conjugated secondary were incubated with the blots to detect GBA2 (rabbit polyclonal antibodies) and Flag-tag primarily (rabbit monoclonal antibodies) and β-actin (mouse monoclonal antibodies) antibodies, respectively. After washing 3 times with PBST, the blots were developed with Luminata Forte Western HRP Substrate, according to the manufacturer’s instructions (Merck).

### Measurement of GBA2 enzyme activity on MUG

GBA2 enzyme activity was assayed as described elsewhere (8, 20, 21). Samples were pre-incubated with or without CBE, followed by incubated with the 4-methylumbelliferyl-β-D-glucoside (4MUG) substrate, 3.5 mM final concentration (Sigma-Aldrich) at pH 5.8 and 37 °C for 30 min.The reactions were terminated by adding 200 μL 1 M of glycine, pH 10.6, then the fluorescent signal was measured in a fluorescence microplate reader with excitation at 355 nm and emission at 460 nm.

### Lipid extraction and lipid measurement

Cell pellets and extraction blank were freeze dried overnight and stored in −80 °C until use. The samples were subjected to monophasic methanol/chloroform/water lipid extraction, as previously described (22, 23). The supernatants were transferred to 2.0 ml glass vials, and stored at −80 °C until further use. Ten microliters of lipid extracts were evaporated, then washed with 10 mM NH_4_HCO_3_, followed by reconstitution in 40 μl of isopropanol: methanol: chloroform (4:2:1, v: v: v, containing 20 mM ammonium formate. The solutions were then placed into the wells of an Eppendorf twin-tec 96-well PCR plate, and the plate was sealed with sealing tape. Samples were then aspirated via direct infusion nanoESI into an ultra high resolution / accurate mass Thermo Scientific model Orbitrap Fusion™ Lumos™ Tribrid™ mass spectrometer with an Advion Triversa Nanomate nESI source (Advion, Ithaca, NY, USA), operating with a spray voltage of 1.2 kV in positive mode and 1.4 kV in negative mode, and a gas pressure of 0.3 psi, as described (24, 25). For the mass spectrometer, the ion transfer capillary temperature was set to 150 °C, the RF-value to 10%, and the AGC target to 2×10^5^. Spectra were acquired at a mass resolving power at 500,000 (at 200 m/z). Peaks corresponding to the target analytes and internal standards (ISs) were identified by automated peak finding and then assigned at the ‘sum composition’ level of annotation using a developmental version of Lipid Search 5. 0α software (Mitsui Knowledge Industry (MKI), Tokyo, Japan, and Thermo Fisher Scientific) by searching against an accurate mass-based, user-defined database. The search parameters were: Parent (noise) Threshold: 150: Parent (mass) tolerance: 1.5 ppm, Correlation threshold (%): 0.3, Isotope threshold (%): 0.1, Max isotope number: 1 (i.e., including the M+1 peak). Peak detection was set to profile and merge mode to average. The internal standards were used to calibrate the mass spectra prior to database searching. Semi-quantitative analysis of identified endogenous lipids was performed by comparison of their peak areas to the peak areas of the relevant internal standards (The SM internal standard was used for sphingomyelin species and the Cer internal standard was used for ceramide and hexosylceramide species), and by further normalizing to total protein (μg). Note that at the level of annotation acheived using this method, glucosylceramide (GlcCer) and galactosylceramide (GalCer) lipids may not be not differentiated from each other, so are collectively assigned here as hexosylceramides.

### Sequence analysis

Nine GBA2 isoforms listed in the National Center for Biotechnology Information (NCBI) Gene database entry Locus 57704, including isoform 1, isoform 2, isoformX1, isoformX2, isoformX3, isoformX4, isoformX6, isoformX7 and isoformX8, were aligned in MEGA10. Homology modeling was done in the SWISS-MODEL server (https://swissmodel.expasy.org) (26) with the *Tx*GH116 β-glucosidase structure as template (27), and models were visualized in PyMOL (Schrödinger LLC, Portland, OR, USA).

### Statistical analysis

Results are expressed as the mean ± SD. For statistical analysis, the quantitative data were analyzed by 1-way ANOVA and Tukey’s post-hoc test for multiple comparisons via GraphPad Prism 5.0 software. The mean differences were considered significant at p<0.05.

## RESULTS

### Sequence analysis of GBA2 isoform

Thirteen GBA2 isoforms are listed in the National Center for Biotechnology Information (NCBI) Gene database Gene ID 57704 entry. Experimentally determined cDNA sequences for only two of these, isoform 1 and isoform X1, are found in the database, while the rest are supported by high throughput mRNA sequencing data (RNASeq). We have evaluated the functionality of the nine of these GBA2 isoforms that cover most of the gene and do not contain other start codons before their putative start codons by analyzing their effects on the putative structure, and by expression of the corresponding cDNA in COS-7 cells.

GBA2 isoform protein sequences are aligned in Fig. 1. Isoform 1 is the well-characterized standard form of GBA2, while isoform X1, for which a cDNA has been isolated from substantia nigra (AK295967.1), differs only by the insertion of 6 amino acid residues in the N-terminal domain. Isoforms X2 and X4 share this same insertion. Isoforms 2, X4, X6 and X8 have an alternative C-terminus, which is shorter than that found in isoform 1. Isoforms X2, X3 and X6 are missing 22 amino acid residues that contribute to two helices and a loop around the active site in the model of the human GBA2 structure (Supplemental Fig. 1) (27). Isoforms X7 and X8 are missing 79 amino acid residues, which comprise 5 β-strands in the N-terminal domain. Since all of these isoform differences could potentially lead to activity differences, we tested their activities in COS-7 cells.

**Figure 1:**
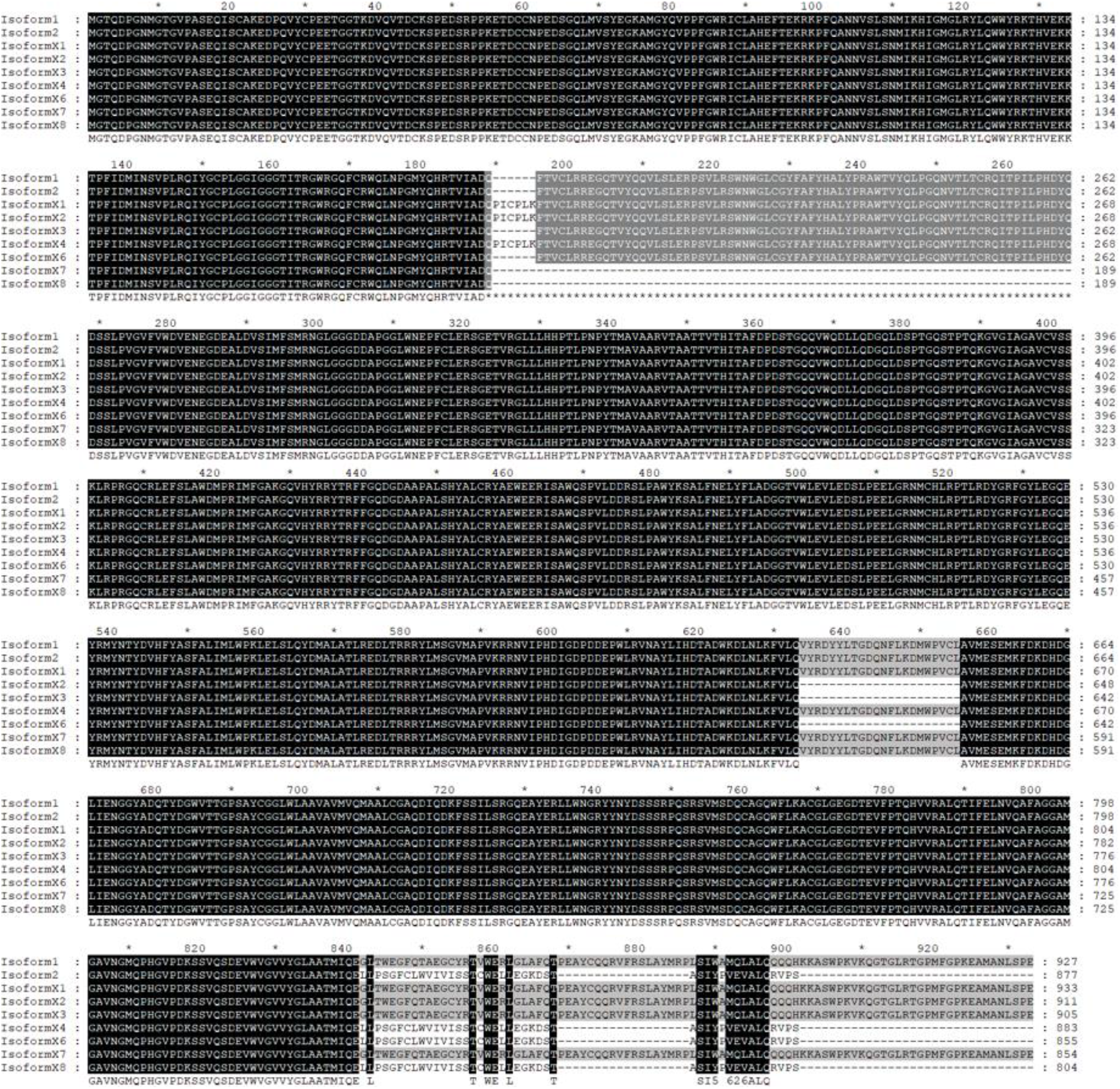
Protein sequence alignment of human GBA2 isoforms.

### GBA2 isoform expression and activity of human GBA2 isoforms in transfected COS-7 cells

Expression analysis by qRT-PCR confirmed that mRNA levels of all human GBA2 isoforms were not significantly different in the same incubation time, while they were significantly decreased at 72 h compared to 48 h (Fig. 2A). Then, we confirmed the expression of each human GBA2 isoform by western blotting using anti-GBA2 and anti-FLAG-tag antibodies (Fig. 2B and 2C), which showed that all human GBA2 isoforms were expressed at the protein level at both 48 h and 72 h. In order to identify which human GBA2 isoforms were active, activity against the fluorescent substrate 4-methylumbelliferyl β-D-glucoside (4MUG) was measured in lysates from transfected COS-7 cells (at 48 and 72 h post-transfection). CBE was added as a GBA inhibitor to one set of assays, although it also exhibits weak inhibition of GBA2 (28).

**Figure 2:**
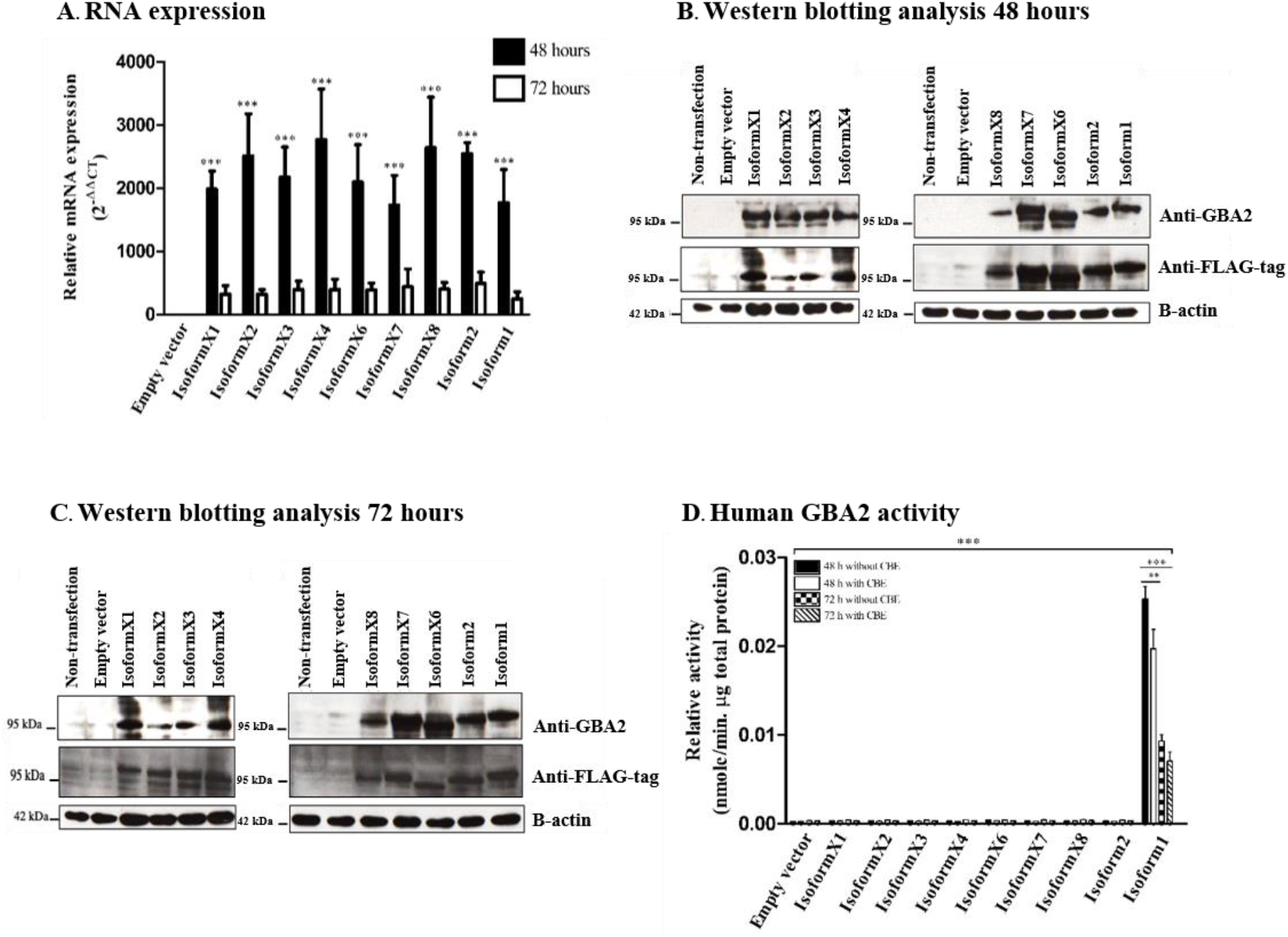
Expression of human GBA2 RNA, protein and activity in COS-7 cells transfected with human GBA2 isoforms. The 9 human GBA2 isoforms were expressed in COS-7 cells, and RNA and protein were extracted at 48 and 72 h. Cell homogenates were also incubated with 4-methylumbelliferyl β-glucoside substrate in reaction buffer to determine β-glucosidase activity. **(A)** RNA expression level of human GBA2 isoforms in transfected COS-7 cells. **(B)** and (**C)** Protein expression of human GBA2 in transfected COS-7 cells detected with anti-human GBA2 and anti-FLAG-tag, respectively at 48 h and 72 h post-transfection, respectively. **(D)** β-Glucosidase activity in extracts of cells expressing human GBA2. Result are representative of three independent biological replicates, *p<0.05 and ***p<0.01.

After 48 h of transfection, we found that human GBA2 activity of cells transformed with isoform 1 was significantly higher than cells transfected with empty vector as a negative control, while cells with other isoforms were similar to the negative control. Seventy-two hours after transfection, the human GBA2 activity of cells transfected with isoform 1 was significantly lower than after 48 h (Fig. 2D). Furthermore, GBA2 activity decreased about 25% upon addition of CBE, confirming that most of the activity resulted from GBA2 rather than endogenous GBA activity. Thus, all human GBA2 isoforms were expressed but only isoform 1 clearly hydrolyzed the MUG substrate. These results suggest that none of the deletion or insertions shown in the protein sequence alignment in Fig.1 and structural model (supplemental Fig. 1) can be accepted and still form an active 4MUG hydrolase.

### Analysis of sphingolipid levels in COS-7 cells overexpressing GBA2 isoforms

The above results confirmed the GBA2 isoform 1 expressed in COS-7 cells can hydrolyze the synthetic 4MUG substrate, while other isoforms showed little or no activity with this substrate. However, this did not indicate whether human GBA2 isoform 1 or the other isoforms can act on natural glucosylceramide substrates in the cells. Therefore, the lipid levels in COS-7 cells expressing the isoforms at 48 h and 72 h post-transformation were determined by UHRAMS analysis. The levels of total sphingolipid species identified at the 48 h time point, including total sphingolipid, total ceramide, total hexosylceramide and total sphingomyelin, are shown in Fig. 3, while the sphingolipid species identified at the 72 h time point are shown in supplemental Fig. 2. Total ceramide was not significantly increased by GBA2 isoform 1 overexpression, while total hexosylceramide decreased, albeit not by a significant amount, and total sphingolipid and sphingomyelin underwent an insignificant increase in cells expressing GBA2 isoform 1 compared to the control. The heat map of the sphingolipid concentration Z-scores at 48 h post-transfection in Fig. 4A shows the ceramide (Cer), hexosyl ceramide (HexCer) and sphingomyelin lipids that were identified in all conditions. The identified ceramides (Cer) had total numbers of carbons of C32-42, including the sphingoid base and long chain fatty acid, while identified hexosyl ceramides (HexCer) had lipid components of C38-42, and identified sphingomyelins (SM) included species with C32-44. Interestingly, clustering of the cell extracts based on their sphingolipid compositions identified GBA2 isoform 1 as the outgroup, which is what would be expected if it is the only isoform with significant activity on the sphingolipids.

**Figure 3:**
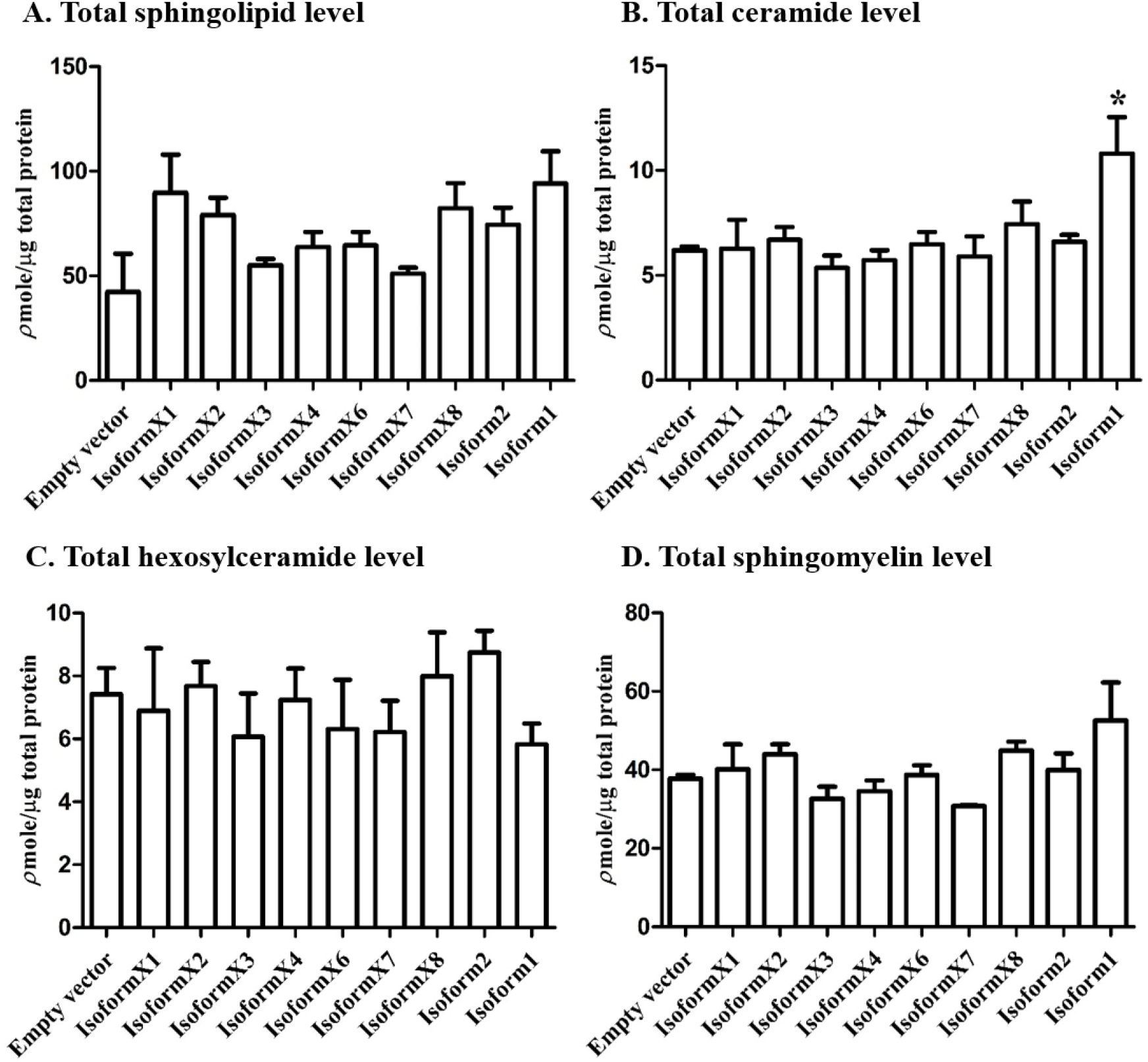
Profiles of total sphingolipid, total ceramides, total hexosylceramide and total sphingomyelin in COS-7 cells transfected with the 9 human GBA2 isoforms 48 hours post-transfection. All species detected for each sample were included in the sum, regardless of whether they were found in other samples or not. Data are expressed as mean of three independent replicates ± SD, *p<0.05.

**Figure 4:**
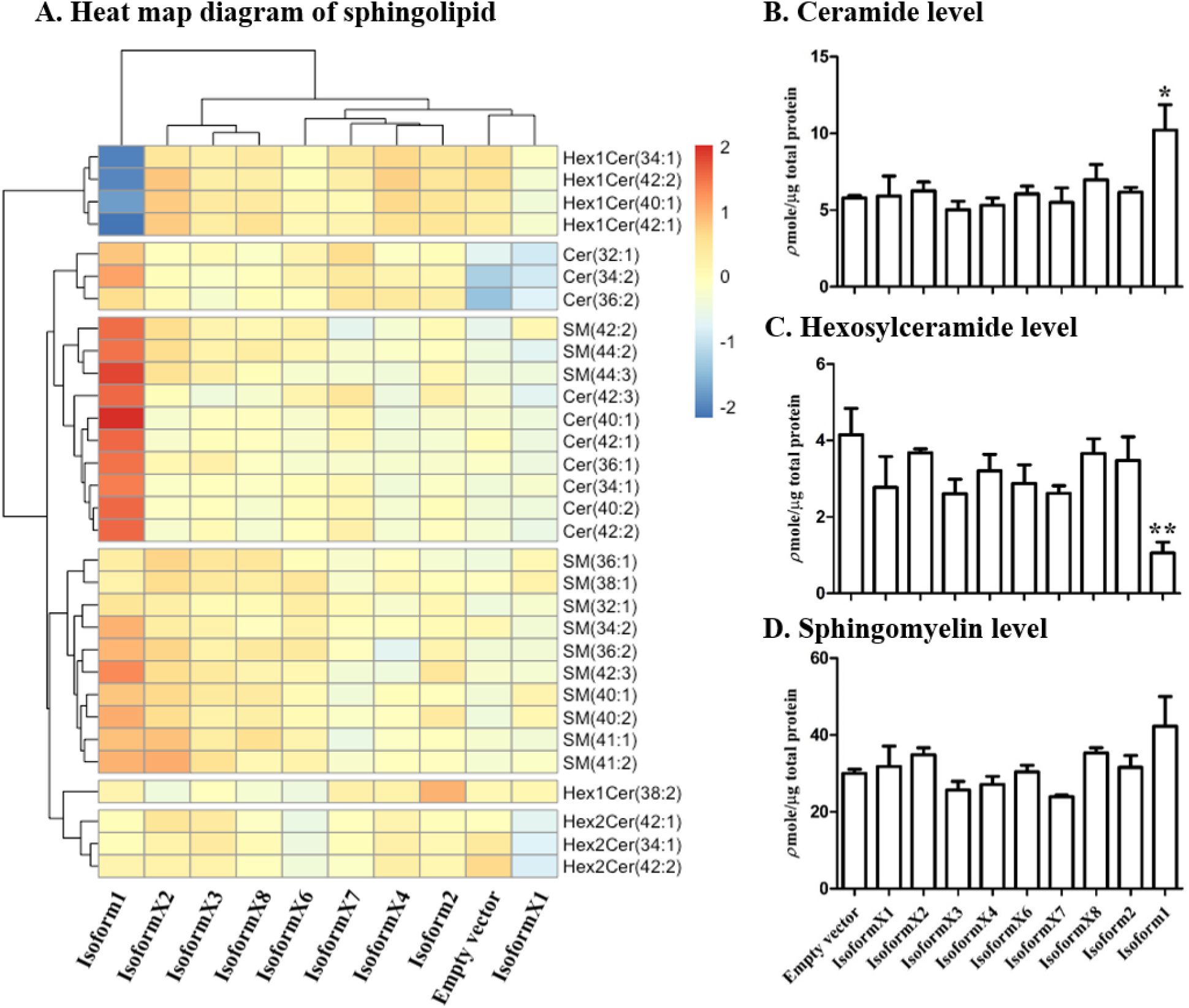
Relative sphingolipid levels in cells expressing respective human GBA2 isoforms. **(A)** The heat map illustrates Z-score differences from mean of sphingolipid in cells expressing the human GBA2 isoforms and control at 48 h after transfection, while the cluster maps illustrate the similarities of the patterns. The z-scores are color-coded from blue (lower than average for that lipid species) to red (higher than average for that lipid species). **(B)** Ceramide (34: 1, 36: 1, 40: 1, 40: 2, 42: 1, 42: 2 and 42: 3), **(C)** Hexosylceramide (34:1, 40:1, 42:1 and 42:2), and **(D)** Sphingomyelin (36:2, 40:1, 40:2, 41:1, 41:2, 42:1, 42: 2, 42: 3, 44: 2 and 44: 3), expressed as bar graphs. Values are means of three independent biological replicates, *p<0.05 and **p<0.01.

In the extract from cells transformed with isoform 1, the mono-hexosylceramide species Hex1Cer(34:1), Hex1Cer(40:1), Hex1Cer(42:1) and Hex1Cer(42:2) showed low concentrations (low Z-scores), while ceramides Cer 34: 1, 40: 1, 42: 1 and 42: 2 were found at high levels relative to the other conditions. The heat map also indicated that SM species 34:2, 36:2, 40:2, 42:1, 42:2, 42:3, 44:2, and 44:3 were higher in isoform 1-expressing cell extracts than in those of other cells. However, other SM species, such as 32:1, 36:1, and 38:1, were at similar or lower levels in isoform 1 expression cell extracts compared to those of other cells. This suggests that GBA2 isoform 1 hydrolyzes GlcCer to release Cer and glucose, resulting in lower hexosylceramide and higher free ceramide levels, while other isoforms did not significantly affect these levels, as emphasized by combining all species of each lipid class in the bar graphs in Fig. 4B and 4C. In contrast, the levels of Hex1Cer(38: 2), and the di-hexosyl (i. e., lactosyl) ceramide species Hex2Cer(34:1), Hex2Cer(42:1) and Hex2Cer(42:2) were similar to or slightly higher than those in the control upon overexpression of GBA2 isoform 1. This suggests that depletion of the glucosylceramide did not have a significant effect on dihexosyl ceramide levels. The differences between SM levels in control cells and cells expressing isoform 1 were not significant, as shown in Fig. 4D. In contrast to isoform 1, the other isoforms did not induce significant changes in cellular sphingolipid levels. Thus, these results confirmed that GBA2 isoform 1 is the only active isoform and indicate that Hex1Cer isoforms 34:1, 40:1, 42: 1 and 42: 2 all appear to be human GBA2 substrates. However, the heat map of the sphingolipid concentration Z-scores at 72 h post-transfection was not significantly different in all conditions, as shown in supplemental Fig. 3, suggesting that other factors had more influence on the sphingolipid levels than expression of GBA2 as its activity decreased.

### Analysis of sphingolipid ratios related to the direction of sphingolipid metabolic flow

To generate a more sensitive parameter for the movement of ceramides from glucosylceramides to other species upon overexpression of GBA2, the ratios of Cer to HexCer and SM to HexCer were calculated. HexCer and Cer with the same ceramide masses that were detected included 34: 1, 40: 1, 42: 1 and 42: 2 species, while of these SM species were detected only for 40: 1 42: 1 and 42: 2. The expression of human GBA2 isoform 1 in COS-7 cells resulted in an increased ratio (of 26) for the Cer to Hex1Cer (34:1, 40:1, 42:1 and 42:2) compared to a ratio of 5.7 for the empty vector control (Fig. 5A). However, expression of the other isoforms did not significantly change these ratios at 48 h, as seen in Fig. 5A. The ratio of SM to HexCer (40: 1, 40: 2 and 42: 2) was also significantly higher in GBA2 isoform 1 transfected COS-7 cells, compared to other isoforms and to control (Fig. 5B). This evidence suggests that GBA2 activity contributes to the conversion of sphingolipid from GlcCer to Cer, and that some of the released Cer may be subsequently converted to SM.

**Figure 5:**
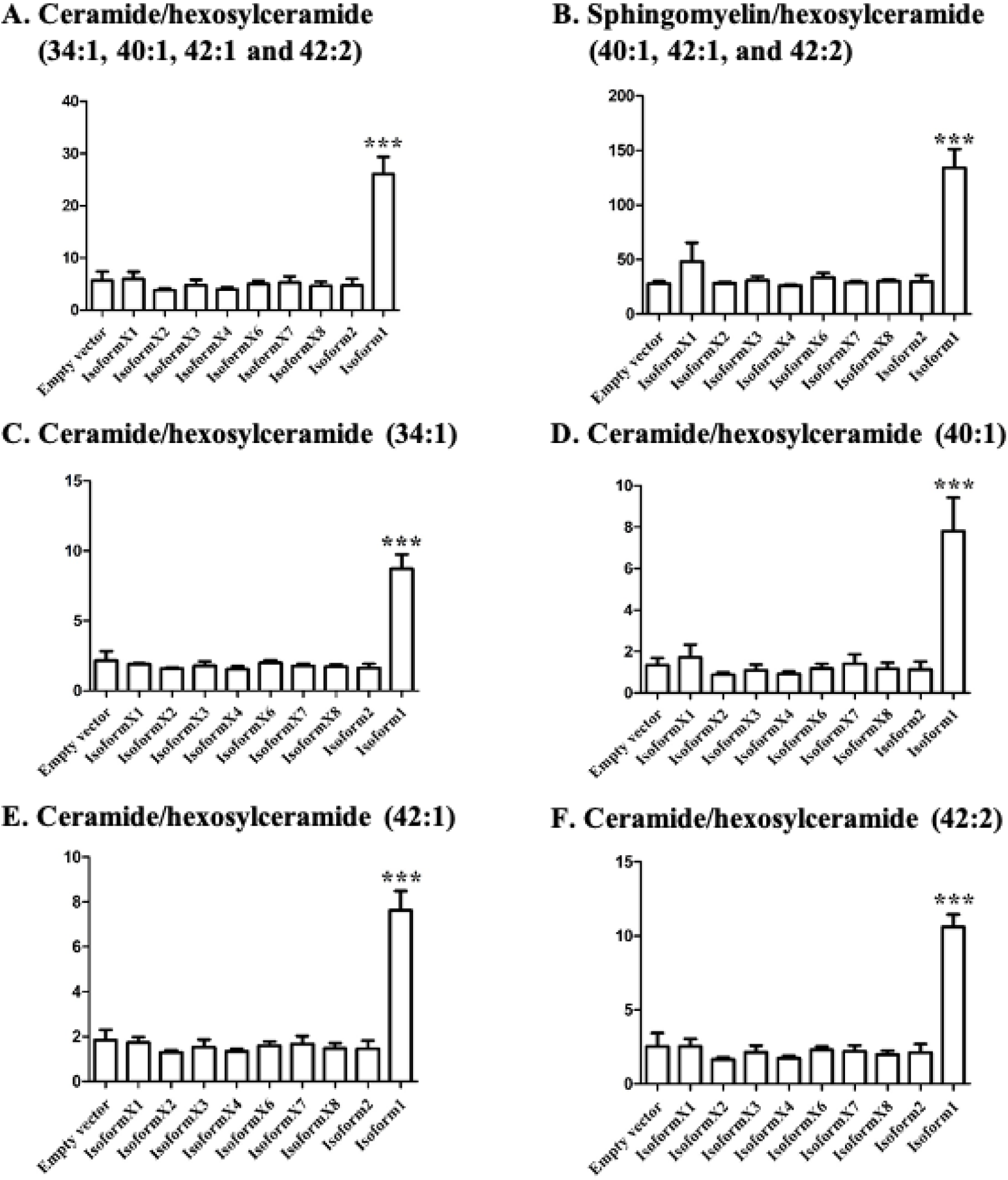
Effect of GBA2 isoforms on ceramide/hexosylceramide ratios of specific lipid species. Sphingolipid ratio **(A)** Ceramide/hexosylceramide (34:1, 40:1, 42:1 and 42:2), **(B)** sphingolyelin/hexosylceramide (40:1 42:1 and 42:2) and ceramide/hexosylceramide ratios for 34:1, 40:1 42:1 and 42:2 are shown separately in **(C), (D), (E),** and **(F)**, respectively. Data are expressed as mean of three independent replicates ± SD, ***p<0.01.

Since a GBA2 isoform could act on one or a subset of the glucosylceramide species, the change in intracellular sphingolipid ratio was analyzed for each ceramide/hexosylceramide pair, including HexCer to Cer (34:1) in Fig. 5C, HexCer to Cer (40:1) in Fig. 5D, HexCer to Cer (42:1) in Fig. 5E, and HexCer to Cer (42:2) in Fig. 5F. The ratios of Cer to HexCer were increased for cells transfected with GBA2 isoform 1, but not with the other isoforms. At 72 h, the ratio of Cer to HexCer and SM to HexCer were not significantly different from control, as shown in supplemental Fig. 4. These results suggest that each pair represents a GBA2 isoform 1 substrate and product, while the other isoforms show no obvious activity toward any of them.

### Analysis of total lipid composition

The total lipid distribution among the 3 lipid classes, including sphingolipids, glycerophospholipids and glycerolipids did not change significantly in cells overexpressing human GBA2 isoform 1 compared to control as shown in Fig. 6 (48 h post-transfection) and supplemental Fig. 5 (72 h post-transfection). Consideration of the amounts of individual species of each lipid class in Fig. 6B, 6C and 6D shows that sphingolipids and glycerolipids appeared to increase slightly, while the average amounts of phosphoglycerolipids did not change significantly. These results suggest that overall lipid homeostasis was not disturbed by human GBA2 overexpression.

**Figure 6:**
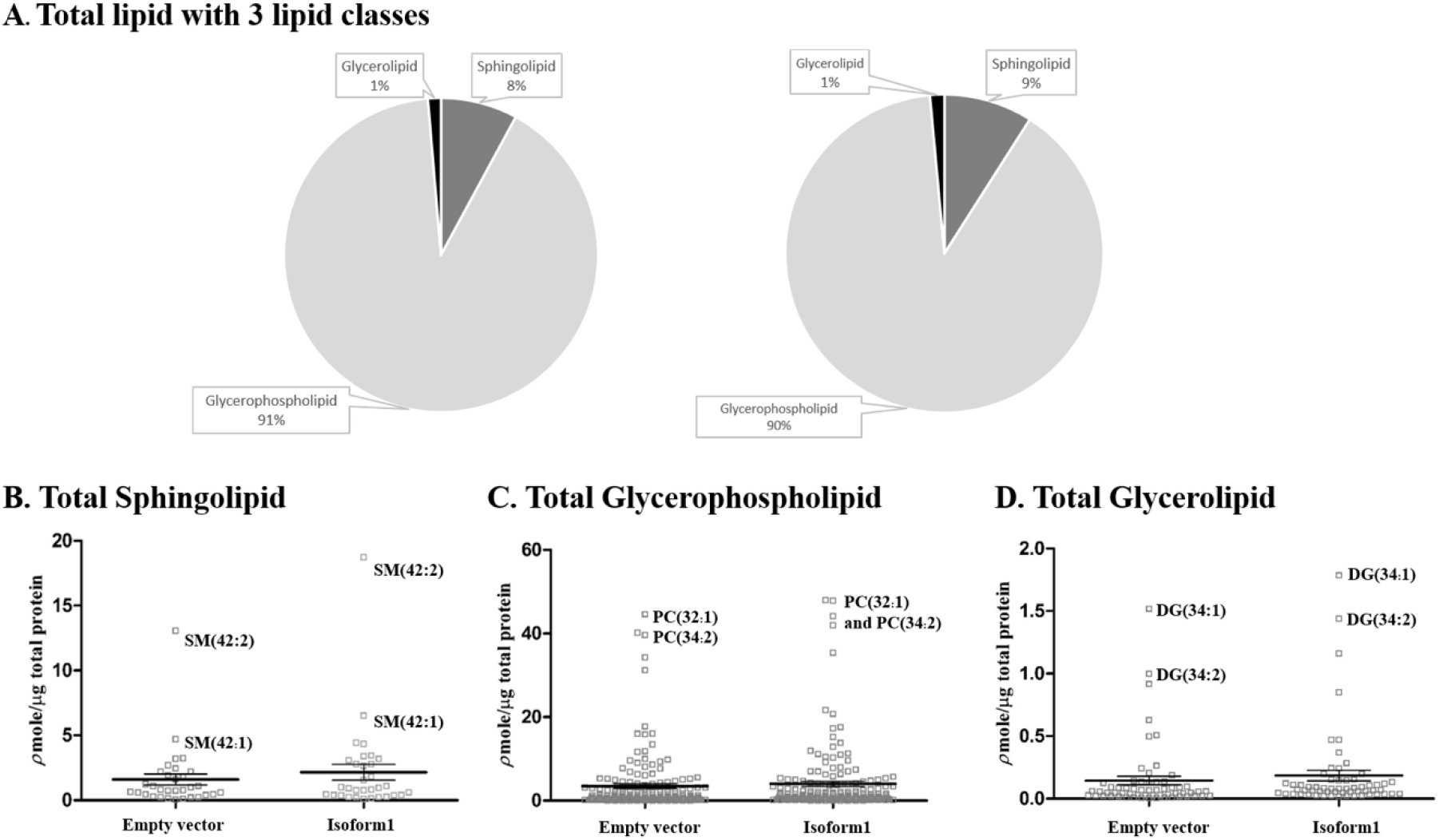
Relative levels of sphingolipids, glycerophospholipids and glycerolipids in COS-7 cells transfected with empty vector and vector for human GBA2 isoform 1 for 48 hours. **(A)** Relative amounts of 3 classes of lipid species. Levels of specific sphingolipid **(B)** glycerophospholipid **(C**) and glycerolipid **(D)** species and average values in control and cells expressing GBA2 isoform 1 are illustrated as parallel dot plots. Amounts were determined by mass spectrometry analysis of COS-7 cell lipid extracts. Result are representative of three independent replicates, *p<0.05.

### Analysis of glycerophospholipids/glycerolipids involved in sphingolipid metabolism

In the previous analysis, Cer was increased in response to overexpression of GBA2 glucosylceramidase, and this appeared to cause an increase in some SM levels as well. Since SM synthase transfers a phosphocholine group from phosphotidylcholine (PC) to Cer to produce SM and diacylglycerol (DAG) (29, 30), we analyzed the ratio of total DAG to PC, at 48 h post-transfection in Fig. 7A. DAG and PC species with fatty acyl components of 34: 1, 40: 2, 40: 3, 40: 5 and 40: 6 (number of carbons: double bonds) were detected in both control and GBA2-overexpressing cell extracts. The ratio of DAG to PC was increased for the total of all of these species and for each species independently in cells overexpressing GBA2 isoform 1 compared to control. However, this increase was only significant for DAG to PC with the fatty acyl component 34:1, while no significant differences were observed at 72 h (supplemental Fig. 6). In comparison, no significant differences were observed in the ratios for DAG to PE and PI (supplemental Fig. 7 and 8).

**Figure 7:**
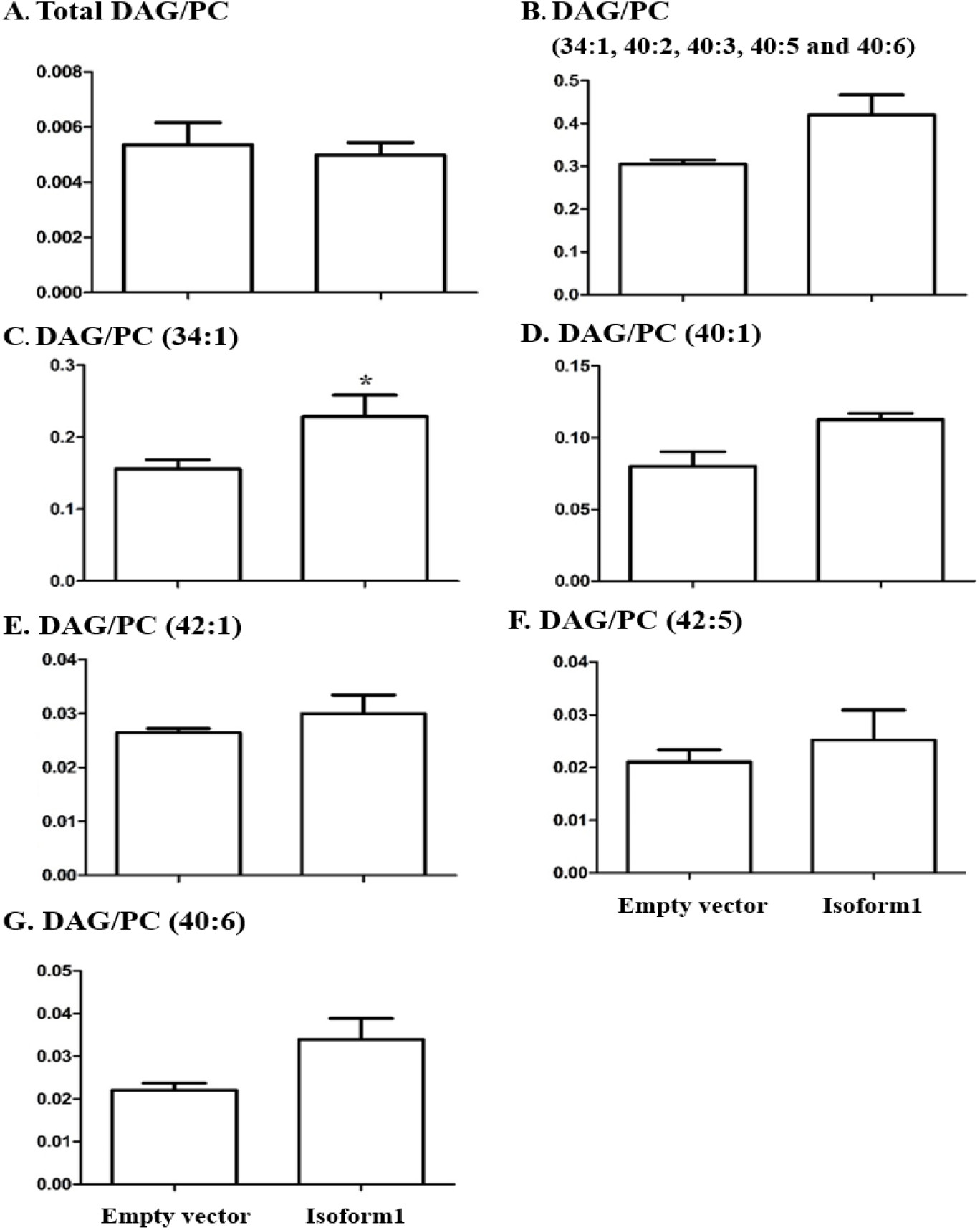
Ratios of levels of diacylglycerol to phosphatidylcholine in COS-7 cells transfected with control vector and GBA2 isoform 1 expression vector for 48 hours. The ratios of total diacylglycerol (DAG) to total phosphatidylcholine (PC) are shown in **(A**). Total DAG/PC for those lipid species found in both lipid classes 34: 1, 40: 2, 40: 3, 40: 5 and 40: 6) are shown in (**B**). The individual DAG/PC ratios for 34:1, 40:1, 42:1 and 42:2 are shown in (**C), (D), (E), (F)** and (**G**), respectively. Amounts in lipid extracts of COS-7 cells transfected with empty vector and human GBA2 isoform 1 expression vector for 48 h were determined by mass spectrometry analysis. Means and standard deviations of ratios in three independent biological replicates are represented, *p<0.05.

## DISCUSSION

GBA2 deficiency is responsible for a heterologous group of ataxias, including hereditary spastic paraplegia, autosomal recessive cerebellar ataxia with spasticity and Marinesco-Sjogren-like syndrome (31–33). The molecular basis of how GBA2 deficiency leads to these syndromes and the basis for their heterogeneity are not well understood. We considered the possibility that some splice isoforms might show differential tissue expression and certain mutations might affect some of these more than others. Although previous papers only considered GBA2 isoform 1 (8, 16, 31, 32, 34), twelve other isoforms are predicted from RNA sequencing (RNA Seq) data in the NCBI database, with some resulting in a seemingly mild change in the noncatalytic domain. However, when we expressed the nine human GBA2 isoforms with the most complete sequences, only isoform 1 had activity toward MUG and caused a significant change in cellular sphingolipids. Surprisingly, isoformX1, which has an insertion of 6 amino acid between residues 189 and 190 in a β-strand of the N-terminal non-catalytic domain compared to isoform 1, showed no significant activity, indicating that the structural integrity of the N-terminal domain is also critical to the activity. It has recently been demonstrated that GBA2 forms oligomers, which may be necessary for its function (35), so disruption of oligomerization could be one explanation for this defect.

Although previous papers have expressed GBA2 in cells, including studying the effects of mutations (8, 35), these generally only looked at the activity on the synthetic substrate MUG. In other cases, the effect of deficiency of GBA2 on sphingolipid levels was explored in animals (16, 34, 36). For instance, glucosylceramides with various lipid components were found to build up in testis and dermal fibroblasts of homozygous GBA2 knockout mice (37). Consistent with those results, overexpression of GBA2 in COS-7 cells in this work resulted in decreased levels of specific HexCer, corresponding to glucosylceramides. In contrast to that previous work in mice, we detected a clear increase in ceramides, while no significant decrease in ceramides was seen in GBA2 deficiency in mouse testes or fibroblasts (36). Similar increases in GlcCer were seen in the brains of Niemann-Pick Type C model mice, when GBA2 was knocked out or inhibited (37). Glucosylceramides were also significantly higher in lymphoblastoid cells from a patient with a homozygous GBA2 mutation compared to control lymphoblastoid cells (20). So, in general, our observations in overexpression of GBA2 on sphingolipids are opposite to those seen in animal and human cells with GBA2 deficiency, as expected, except that we also detected a significant change in ceramide levels.

Upon comparing the general levels of hexosylceramides in Fig. 3, the decrease seen upon overexpression of GBA2 isoform 1 was not significant, partly due to the overlapping presence of both glucosylceramides and galactosylceramides, and species that were detected in some samples and not others. By comparing only species of lipids found as both ceramides and hexosylceramides in all samples in Fig. 4B, we were able to see a significant change. The Fig. 4A heat map showed that HexCer(42: 1), (42: 2), (34:1) and (40:1) increased significantly compared to control, while HexCer(38:2) was similar in cells with overexpressed GBA2 isoform 1 and control. This suggests that HexCer(38: 2) may have a higher fraction galactosylceramides compared to glucosylceramides, while the other 4 HexCer masses represented higher fractions of glucosylceramides that could be hydrolyzed by GBA2. Despite the obvious SM increase seen upon overexpression of GBA2 isoform 1 in the Fig. 4A heat map, the change was not to a significant level. To develop a more sensitive parameter, we evaluated the ratio of ceramide to hexosylceramide for lipid species found in both classes in Fig. 5, and found that the difference upon GBA2 isoform 1 expression was much more highly significant. The ratio of sphingomyelins to hexosylceramides of the same species showed a similar level of significance (Fig. 5A), suggesting that some hexosylceramide hydrolyzed by the overexpressed GBA2 is likely to be converted to sphingomyelin.

Since sphingomyelin synthase transfers phosphocholine from PC to ceramide to make SM and release DAG, we also investigated the ratios of DAG to PC with the same fatty acid masses. As seen in Fig. 7, the ratios all increased for those species detected in both PC and DAG, although only in the case of the 34:1 species (likely corresponding to one oleic acid (18:1) and one palmitic acid (16:0)) was the increase statistically significant. The overall lipid proportions were not disrupted (Fig. 6), and no significant differences in levels of total glycerophospholipids, phosphatidylcholine, phosphatidyl ethanolamine, phosphatidlylinositol and glycerolipids were observed compared to other isoforms and empty vector, as shown in supplemental Fig. 2. The results suggest that overall lipid homeostasis was not generally disrupted by the GBA2 overexpression, but the levels of certain DAG species may have increased, along with Cer and SM, since PC levels are high and unlikely to be significantly affected by their use for SM synthesis. Given the role of DAG in protein kinase C activation and signaling (38), this could be another aspect of GBA2 deficiency or excess GBA2 activity.

Although natural cases of overexpression of GBA2 have not been demonstrated, GBA2 activity is increased by high substrate concentrations when lysosomal GBA is deficient in Gaucher and Niemann-Pick models (28), some symptoms of which are decreased when GBA2 is knocked out or inhibited (34, 37). These symptoms were suggested to be caused by release of sphingosine in the cytoplasm, but our data suggest that changes in ceramide, glucosylceramide and DAG levels should also be considered.

In conclusion, our work has demonstrated that among the possible isoforms predicted from RNA sequencing in human tissues, only GBA2 is likely to affect the cellular lipid levels directly, although we cannot rule out regulatory roles for other isoforms or their RNA molecules. GlcCer and Cer levels are affected most clearly by GBA2 overexpression, but subtle effects on SM and DAG/PC levels were also seen. Given the effects of these lipids on membrane properties and signaling, GBA2 expression levels may have a significant impact on the cell.

## Abbreviations

CDase: ceramidase
CerS: ceramide synthase
Cer: ceramide
CBE: Conduritol-β-epoxide
DAG or DG: diacylglycerol
ER: endoplasmic reticulum
GSL: glycosphingolipid
GBA2: glucosylceramidase 2
GBA: glucosylceramidase
GlcCer: glucosylceramide
GalCer: galactosylceramide
HexCer: hexosylceramide
HexCer(d18: 1/16: 0): hexosylceramide with a C18 sphingosine (d18: 1) and N-acyl group (16: 0)
MG: monoacylglycerol
LPC: Lysophosphatidylcholine
LPE: Lysophosphatidylethanolamine
PA: Phosphatidic acid
PC: phosphatidylcholine
PE: phosphatidylethanolamine
PG: Phosphatidylglycerol
PI: phosphatidylinositol
PS: phosphatidylserine
SL: sphingolipid
SM: sphingomyelin
SMase: sphingomyelinase
SMS: sphingomyelin synthase
Sph: sphingosine
SPT: serine palmitoyltransferase
S1P: sphingosine-1-phosphate
4MUG: 4-Methylumbelliferyl-β-D-glucopyranoside
TG: triacylglycerol

## Acknowledgement / grant support

The authors would like to thank Dr. Vinzenz Hofferek for advice on the mass spectrometry and software analysis. Financial support was provided by Suranaree University of Technology, including the National Research University Project of the Commission on Higher Education, and also from the Australian Research Council to GER (LE160100015). PJ was supported by a Development and Promotion of Science and Technology Talents Project scholarship.

